# An average-case sublinear exact Li and Stephens forward algorithm

**DOI:** 10.1101/322396

**Authors:** Yohei M. Rosen, Benedict J. Paten

## Abstract

Hidden Markov models of haplotype inheritance such as the Li and Stephens model allow for computationally tractable probability calculations using the forward algorithms as long as the representative reference panel used in the model is sufficiently small. Specifically, the monoploid Li and Stephens model and its variants are linear in reference panel size unless heuristic approximations are used. However, sequencing projects numbering in the thousands to hundreds of thousands of individuals are underway, and others numbering in the millions are anticipated.

To make the Li and Stephens forward algorithm for these datasets computationally tractable, we have created a numerically exact version of the algorithm with observed average case 𝒪(*nk*^0.35^) runtime, avoiding any tradeoff between runtime and model complexity. We demonstrate that our approach also provides a succinct data structure for general purpose haplotype data storage. We discuss generalizations of our algorithmic techniques to other hidden Markov models.

**2012 ACM Subject Classification:** Theory of computation ⟶ Streaming, sublinear and near linear time algorithms; Applied computing ⟶ Bioinformatics

**Supplement Material:** https://github.com/yoheirosen/sublinear-Li-Stephens.

**Funding:** This work was supported by the National Human Genome Research Institute of the National Institutes of Health under Award Number 5U54HG007990, the National Heart, Lung, and Blood Institute of the National Institutes of Health under Award Number 1U01HL137183-01, and grants from the W.M. Keck foundation and the Simons Foundation.

**Acknowledgements:** We would like to thank Jordan Eizenga for his helpful discussions throughout the development of this work.

## 1 Introduction

Probabilistic models of haplotypes describe how variation is shared in a population. One application of these models is to calculate the probability *P*(*o*|*H*) of a haplotype *o* given the assumption that it is a member of a population represented by a *reference panel* of haplotypes *H*. This computation has been used in estimating recombination rates [8], a problem of interest in genetics and in medicine. It may also be used to detect errors in genotype calls.

Early approaches to haplotype modeling used coalescent [7] models which were accurate but computationally complex, especially when including recombination. Li and Stephens wrote the foundational computationally tractable haplotype model [8] with recombination. Under their model, the probability *P*(*o*|*H*) can be calculated using the forward algorithm for hidden Markov models. Generalizations of their model have been used for haplotype phasing and genotype imputation. Most of these algorithms [10, 1, 14, 3, 12] use the *forward probabilities* calculated as intermediate values in the forward algorithm.

### 1.1 The Li and Stephens model

Consider a *reference panel H* of *k* haplotypes sampled from some population. Each haplotype *h*_*j*_ ∈ *H* is a sequence (*h*_*j*,1_,…,*h*_*j*,*n*_) of alleles at a contiguous sequence 1,…,*n* of genetic sites. Classically [8], the sites are biallelic, but the model extends to multiallelic sites. [11]

Consider an observed sequence of alleles *o* = (*o*_1_,…,*o*_*n*_) representing another haplotype. The monoploid Li and Stephens model (LS) [8] specifies a probability that *o* is descended from the population represented by *H*. LS can be written as a hidden Markov model wherein the haplotype *o* is assembled by copying (with possible error) consecutive contiguous subsequences of haplotypes *h_j_* ∈ *H*.

#### ▸ Definition 1

(Li and Stephens HMM). Define *x*_*j,i*_ as the event that the allele *o*_*i*_ at site *i* of the haplotype *o* was copied from the allele *h*_*j,i*_ of haplotype *h*_*j*_ ∈ *H*. Take parameters

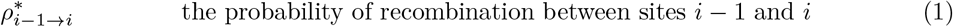

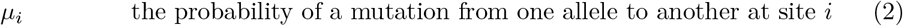

and from them define the transition and recombination probabilities

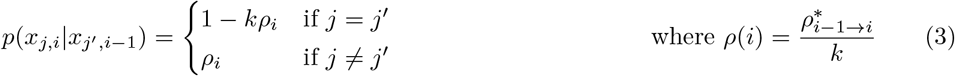

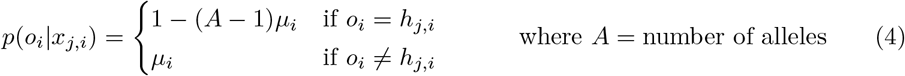

The forward algorithm for hidden Markov models allows calculation of *P*(*o*|*H*) in 𝒪(*nk*^2^) time using an *n* × *k* dynamic programming matrix of *forward states*

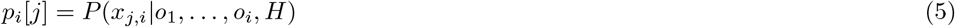

In practice, the Li and Stephens forward algorithm is 𝒪(*nk*). (See §3)

### 1.1.1 Li and Stephens like algorithms for large populations

The 𝒪(*nk*) time complexity of the forward algorithm is intractable for reference panels with large size *k*. The UK Biobank has amassed *k*= 500, 000 array samples. Whole genome sequencing projects, with a denser distribution of sites, are catching up. Major sequencing projects with *k* = 100, 000 or more samples are nearing completion. Others numbering *k* in the millions have been announced. These large population datasets have significant potential benefits: They are statistically likely to more accurately represent population frequencies and those employing genome sequencing can provide phasing information for rare variants.

In order to handle datasets with size *k* even fractions of these sizes, modern haplotype inference algorithms depend on models which are simpler than the Li and Stephens model or which sample subsets of the data. For example, the common tools Eagle-2, Beagle, HAPI-UR and Shapeit-2 and -3 [10, 1, 14, 3, 12] either restrict where recombination can occur, fail to model mutation, model long-range phasing approximately or sample subsets of the reference panel.

Lunter’s “fastLS” algorithm [11] demonstrated that haplotypes models which include all *k* reference panel haplotype could find the Viterbi maximum likelihood path in time sublinear in *k*, using preprocessing to reduce redundant information in the algorithm’s input. However, his techniques do not extend to the forward and forward-backward algorithms.

### 1.2 Our contributions

We have developed an arithmetically exact forward algorithm whose expected time complexity is a function of the expected allele distribution of the reference panel. This expected time complexity proves to be 𝒪(*k*^0.35^) in reference panel size. We have also developed a technique for succinctly representing large panels of haplotypes whose size also scales as a sublinear function of the expected allele distribution.

Our forward algorithm contains three optimizations, all of which might be generalized to other bioinformatics algorithms. In (§2), we rewrite the reference panel as a sparse matrix containing the minimum information necessary to directly infer all allele values. In (§3), we define recurrence relations which are numerically equivalent to the forward algorithm but use minimal arithmetic operations. In (§4), we delay computation of forward states using a lazy evaluation algorithm which benefits from blocks of common sequence. Our methods apply to other models which share certain properties with the monoploid Li and Stephens model.

## 2 Sparse representation of haplotypes

The forward algorithm to calculate the probability *P*(*o*|*H*) takes as input a length *n* vector *o* and a *k* × *n* matrix of haplotypes *H*. Therefore time complexity better than 𝒪(*nk*) is impossible unless there is preprocessing of its input. However, such preprocessing can be amortized over many queries *o*.

### 2.1 Information content of a reference panel

Recall that 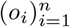 is the allele sequence of the emitted haplotype *o*. (§3) will show that *ϕ*_*i*_(*o*_*i*_, *H*), 1 ≤ *i* ≤ *n* defined below are sufficient data to calculate *P*(*o*|*H*).

#### ▸ Definition 2

The information content *ϕ* of *H* for allele α at site *i* is defined as

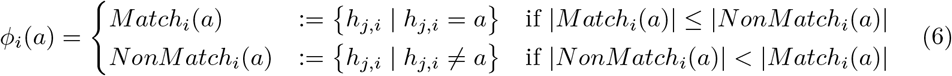

### 2.2 Relation of information content to allele frequency spectrum

Our sparse representation of the haplotype reference panel benefits from the recent finding [6] that the distribution over sites of minor allele frequencies is biased towards low frequencies^2^

We will compute the expected time sum of the information content over all sites assuming first that all sites are biallelic^3^. In the biallelic case *ϕ*_*i*_(·) is always the set of haplotypes displaying the minor allele at site *i* and the distribution of *ϕ*_*i*_(*a*) is the allele frequency spectrum.

#### ▸ Lemma 3

*Let* 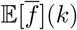 *be the expected mean minor allele frequency for k genotypes. Then*

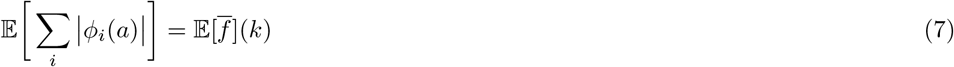

#### ▸ Corollary 4

*if* 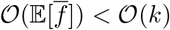, then 𝒪(∑_*i*_|ϕ_*i*_(*a*|) < 𝒪(*k*) *in expected value*.

### 2.3 Implementation

For biallelic sites, we store our *ϕ*_*i*_’s using a length-*n* vector of length |*ϕ*_*i*_| vectors containing the indices *j* of the haplotypes *h*_*j*_ ∈ *ϕ*_*i*_ and a length-*n* vector listing the major allele at each site. (See Figure 1 panel iii) Random access by key *i* to iterators to the first elements of sets *ϕ*_*i*_(*a*) is 𝒪(1) and iteration across these *ϕ*_*i*_(*a*) is linear in the size of *ϕ*_*i*_(*a*). For multiallelic sites, the data structure uses slightly more space but has the same speed guarantees.

**Figure 1.**
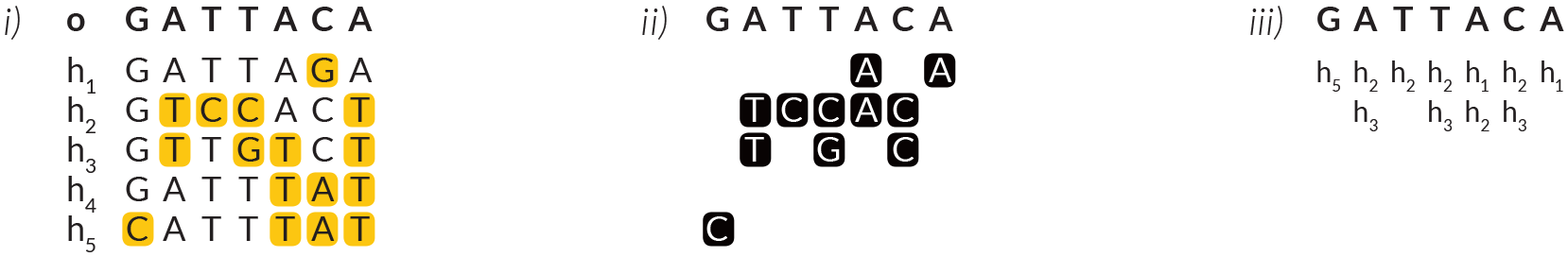
**i)** Reference panel {*h*_1_,…, *h*_5_} with mismatches to haplotype *o* shown in yellow. **ii)** Elements of *ϕ*_*i*_(*o*_*i*_) in black. **iii)** Vectors to encode *ϕ*_*i*_(*o*_*i*_) at each site.

Generating these data structures takes 𝒪(*nk*) time but is embarrassingly parallel in *n*. Our “*.slls” data structure doubles as a succinct haplotype index which could be distributed instead of a large vcf record. A vcf → slls conversion tool is found in our github repository. Adding or rewriting a haplotype is average case constant time per site per haplotype.

## 3 Efficient dynamic programming

We begin with the recurrence relation of the 𝒪(*nk*) Li and Stephens forward algorithm [8]:

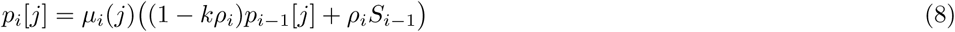

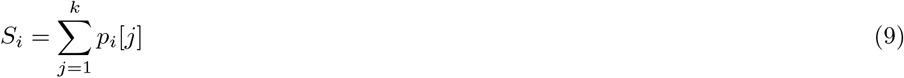

We will reduce the number of summands in (9) and reduce the number indices *j* for which (8) is evaluated, using the **information content** defined in (§2.1).

### ▸ Lemma 5

*The summation (9) is calculable using strictly fewer than k summands.*

**Proof.** Suppose first that μ_*i*_(*j*) = μ for all *j* ≤ *k*. Then

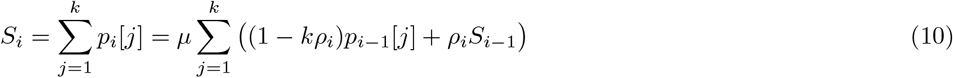

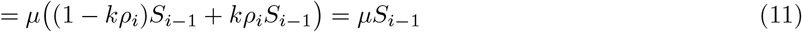

and therefore, relaxing the requirement on μ_*i*_(*j*),

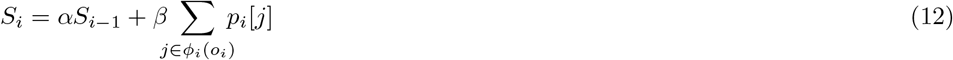

where

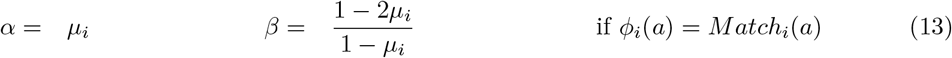

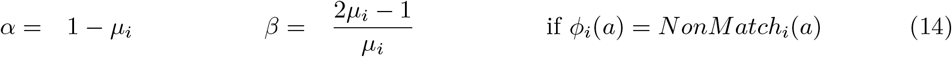

**Figure 2.**
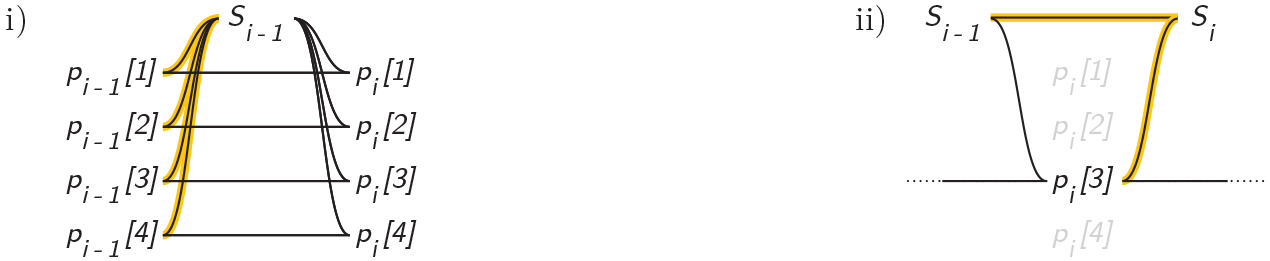
Illustration, with one line per operation, of arithmetic for **i)** conventional 𝒪(*nk*) Li and Stephens recurrence relations **ii)** Our procedure specified in equation (10). Sums in yellow.

### ▸ Lemma 6

*If j* ∉ *ϕ*_*i*_(o_*i*_) *and j* ∉ *ϕ*_*i*−1_(o_*i*−1_), then *S*_*i*_ *can be calculated without knowing p*_*i*−1_[*j*] *and p*_*i*_[*j*], *as can p*_i_[*j*′]*for j*′ ≠ *j*.

**Proof.** By inspection of equation (10).

### ▸ Corollary 7

*The recurrences (9) and the minimum set of recurrences (8) needed to compute (9) can be evaluated in* 𝒪(│*ϕ*│) *time, assuming that p*_*i−*1_[*j*]*have been computed* ⩝ *j*∊ϕ_*i*_(*o*_*i*_).

We address the assumption on prior calculation of the necessary *p*_*i*−1_[*j*]’s in section 4.

### 3.1 Time complexity

Recall that we defined 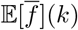 as the expected mean minor allele frequency in a sample of ize *k*. By Corollary 7 the procedure in eq. (10) has expected time complexity 𝒪 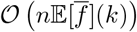.

## 4 Lazy evaluation of dynamic programming rows

Corollary 7 was conditioned on the assumption that specific forward probabilities had already been evaluated. We will describe a second algorithm which performs this task efficiently by avoiding arithmetic which will prove unnecessary at future steps.^4^

### 4.1 Eliminating redundant recurrence evaluations

The recurrence relations (8) are linear maps *r*_*i*_[*j*] : ℝ → ℝ of the form

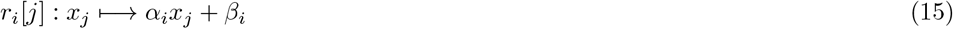

From these, define the linear maps

#### ▸ Definition 8

For any *i*_1_ < *i*_2_, *define the update map*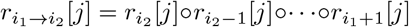 This update map is defined such that 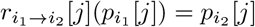.

#### ▸ Lemma 9

*At each i there exist only two unique maps among the r_*i*_[*j*].*

**Proof**. Define 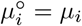 and 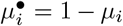 if *ϕ*_*i*_(*a*) = Match_*i*_(*a*) and vice versa otherwise. Then

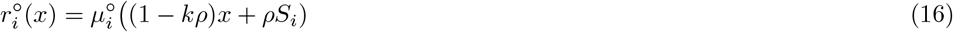

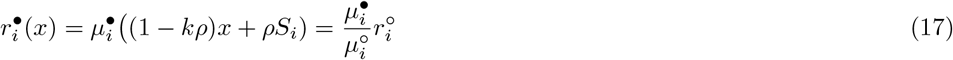

This lemma allows us to rewrite each 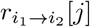 as a binary vector of the form (○,○,•,…,○,•,○).

Our basic delayed evaluation rule is:

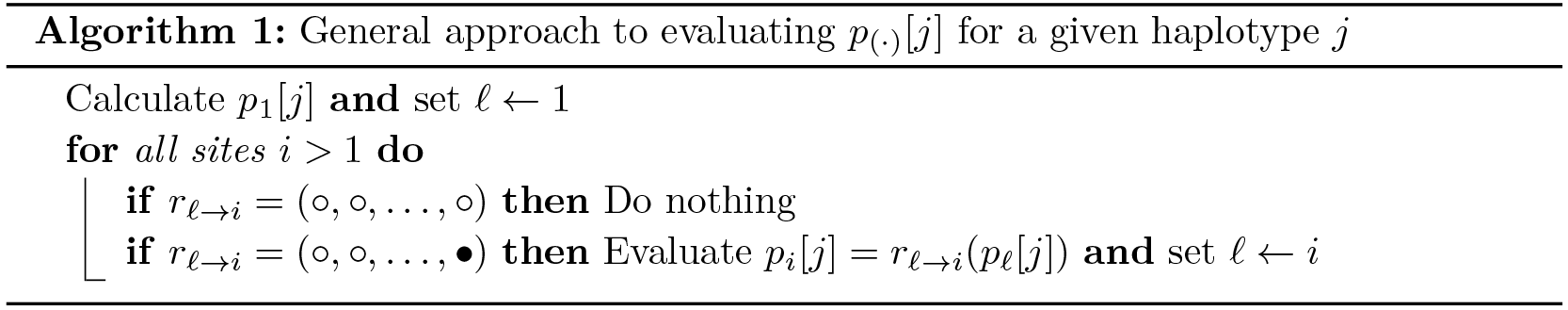

When Algorithm 1 is applied independently to all *h*_*j*_, the aggregate algorithm has 𝒪(*nk*) time complexity, so we will share work between haplotypes *j* using equivalence classes segregated by runs of homology.

### 4.2 Equivalence classes of update map prefixes

Under the conditions of Algorithm 1, if, at step *i*, *p*_(·)_[*j*_1_] and *p*_(·)_[*j*_1_] were both last calculated at the same index ℓ < *i*, then the sequences of *h*_*j*1_ and *h*_*j*2_ are identical between ℓ and *i*, so

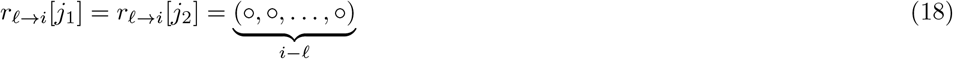

We put *j*_1_ and *j*_2_ into an equivalence class *J*[ℓ] to avoid recalcuating 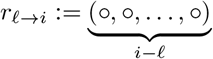

It is also inefficient to construct *r*_ℓ→*i*_ independently for each 1 < ℓ < i. Observe:

#### ▸ Remark

If *J*[ℓ] is empty we need not calculate *r*_ℓ→*i*_.

#### ▸ Lemma 10

If *i*_1_ < *i*_2_ < *i*; then 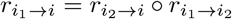

Lemma 10 allows us to calculate intermediate prefixes of the maps *r*_ℓ−*i*_ and extend them at a later time. To make this concrete, suppose that we have an index ◂_ℓ_ where ℓ <◂_ℓ_<*i*.

Then we can evaluate the prefix 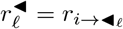 of *r*_ℓ→*i*_ knowing that 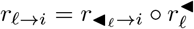 can be evaluated at a later time.

### 4.3 The lazy evaluation algorithm

The data below specifies the state at each step *i* of our lazy evaluation algorithm. The algorithm initialization is described in Algorithm 2 and the recurrence in Algorithm 3.

The maps *j* ↦ J[ℓ];ℓdefined as the index at which *p*_(·)_[*j*] was most recently calculated
The maps 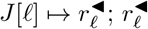 defined as the prefix of *r*_ℓ→*i*_ which was most recently calculated
The maps 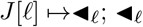 defined as the index up to which a prefix of *r*_ℓ→*i*_ which was most recently calculated
The maps 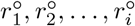

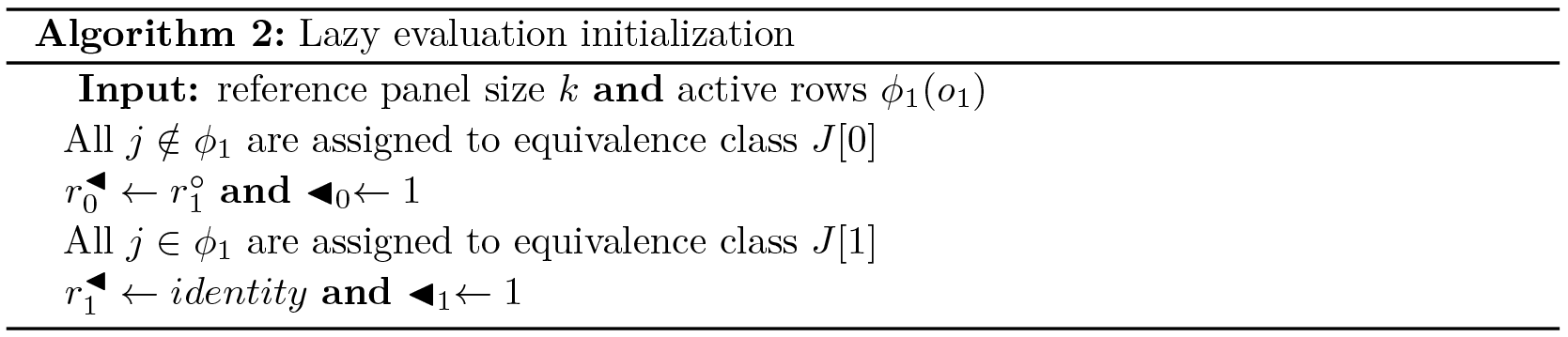

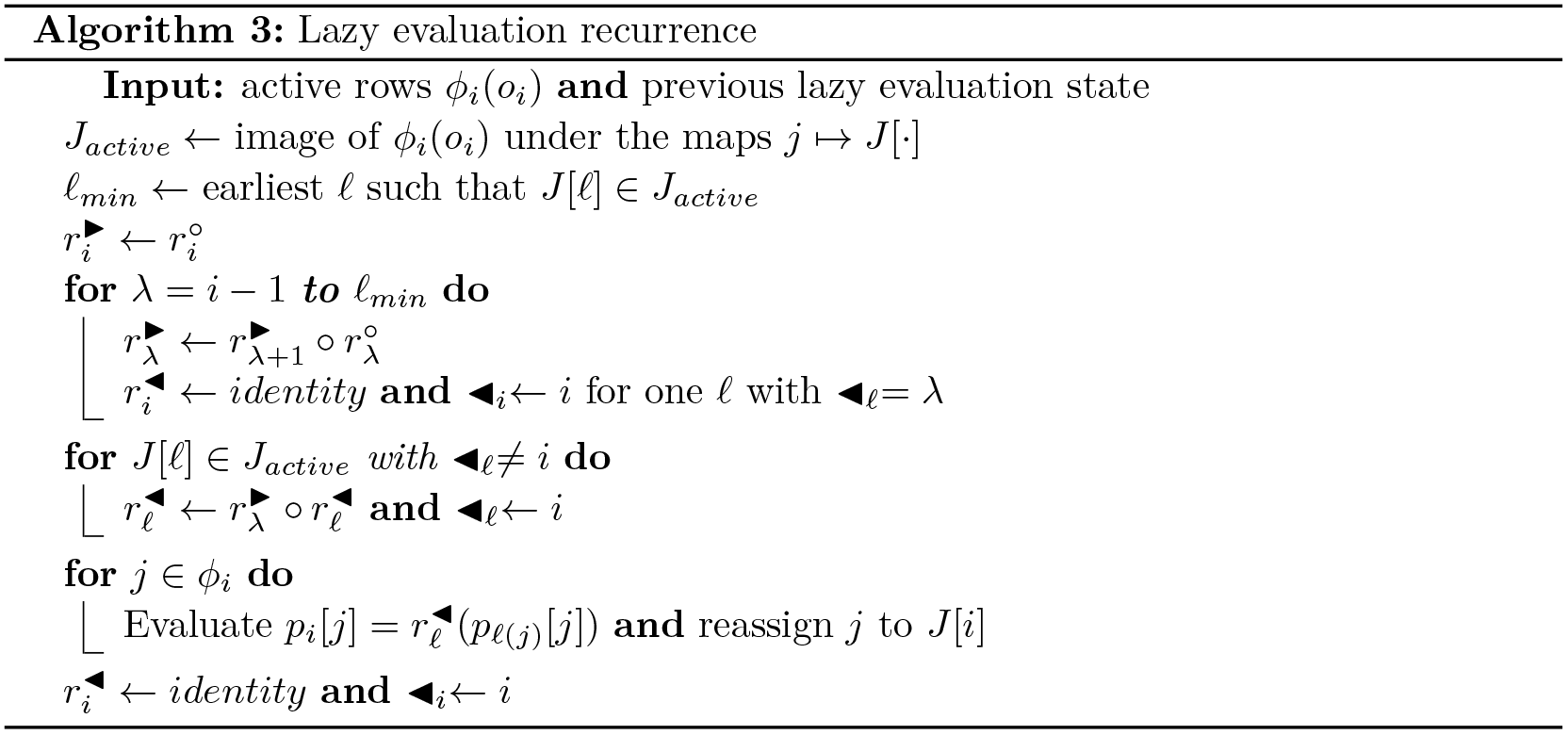

**Figure 3.**
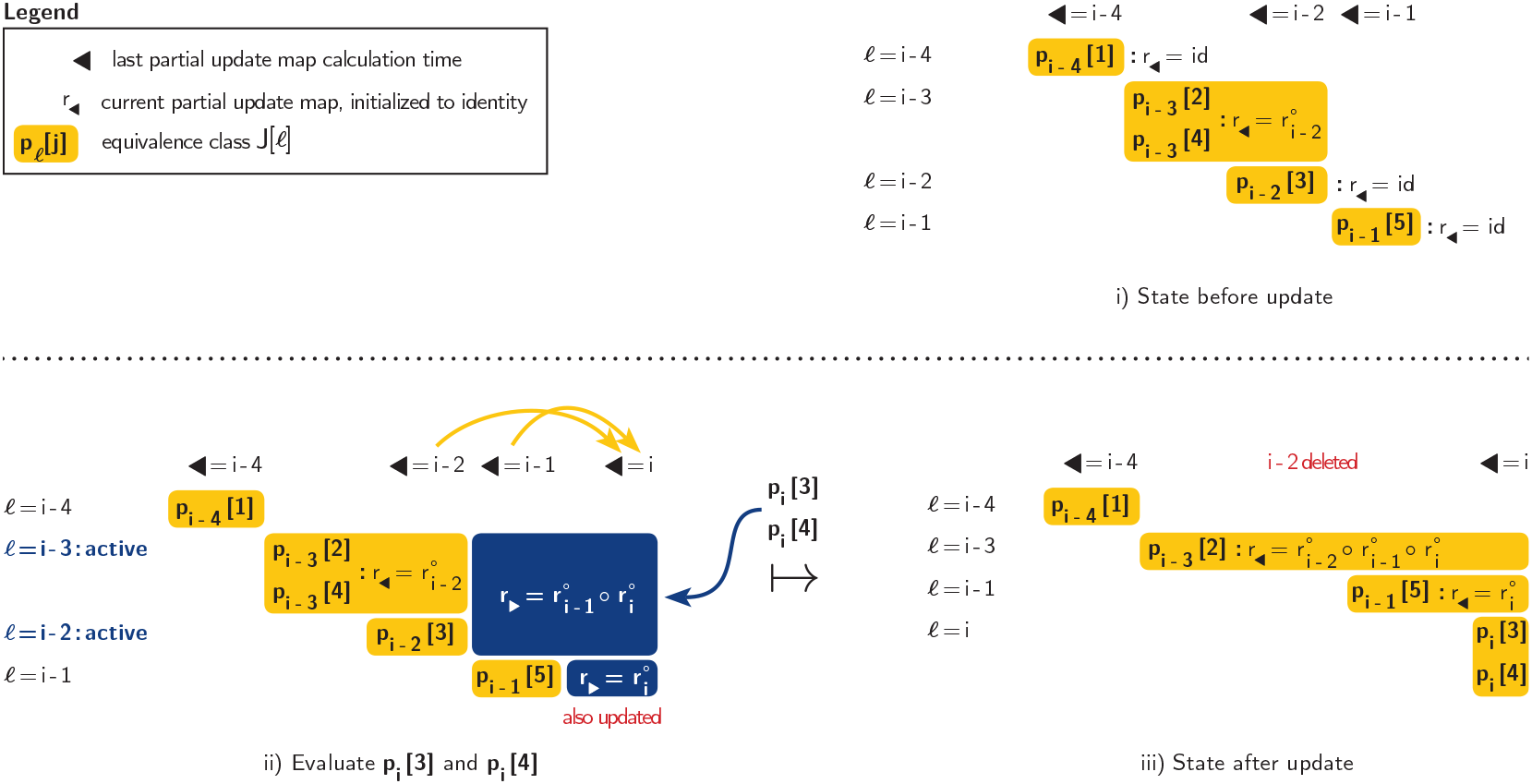
**Top:** Example of lazy evaluation state. **Bottom**: Example of update step as described in Algorithm 3

Calculating a closed form expression for the time complexity of the lazy evaluation algorithm 3 is not straightforward. It is easy to show that it bounded by 𝒪(*nk*), since the first loop is worst-case 𝒪(*k*). However, we find experimentally that asymptotically, this lazy evaluation component does not contribute to overall computational complexity. (See Fig. 6)

## 5 Results

### 5.1 Implementation

Our algorithm was implemented as a C++ library located at https://github.com/yoheirose/sublinear-Li-Stephens Details of 3 will be found there.

We also implemented the linear time monoploid Li and Stephens forward algorithm in C++ as to evaluate it on identical footing. Profiling was performed using a single Intel Xeon X7560 core running at 2.3 GHz on a shared memory machine. Our reference panels *H* were the phased haplotypes from the 1000 Genomes [2] phase 3 vcf records for chromosome 22 and subsamples thereof. Haplotypes *o* were randomly generated simulated descendants.

### 5.2 Minor allele frequency distribution for the 1000 Genomes dataset

We simulated haplotypes *o* of 1,000,000 bp length on chromosome 22 and recorded the sizes of the sets *ϕ*_*i*_(*o*_*i*_) for *k* = 5008. These data produced a mean │*ϕ*_*i*_(*o*_*i*_)│ of 59.9 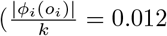 for *k* = 5008.) The distribution (Fig. 4) is skewed toward low frequencies; the minor allele is unique at 71% of sites, and it is below 1% frequency at 92% of sites.

**Figure 4.**
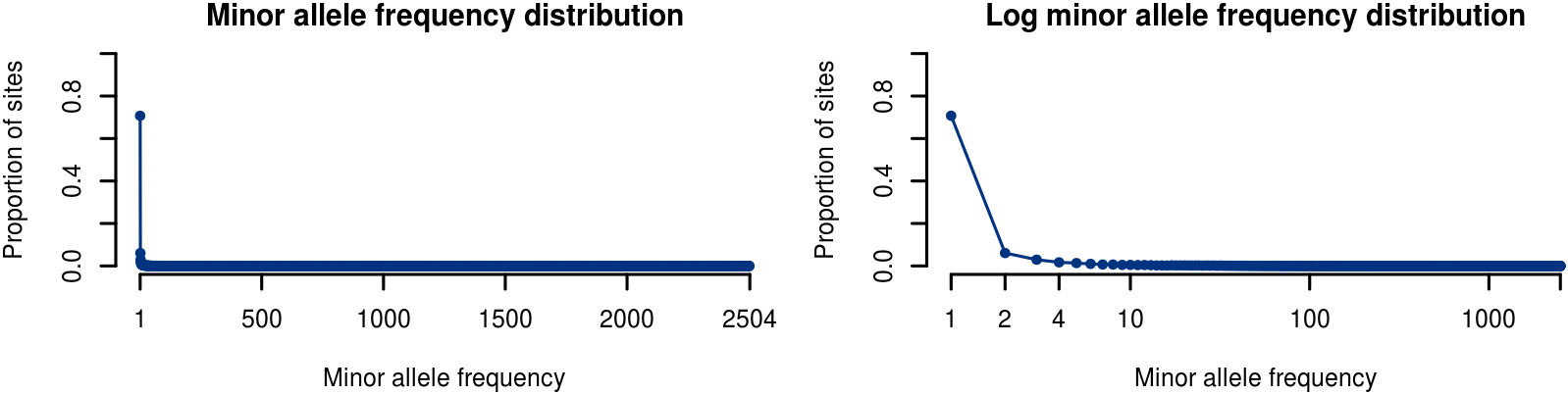
Biallelic site minor allele frequency distribution from 1000 Genomes chromosome 22

### 5.3 Comparison of our algorithm with the linear time forward algorithm

For *k* = 5008, on average, time per site is 37 *μs* for our algorithm and 1308 *μs* for the linear LS algorithm. For the forthcoming 100,000 Genomes Project, these numbers can be extrapolated to 251 *μs* for our algorithm and 260,760 *μs* for the linear LS algorithm.

**Figure 5.**
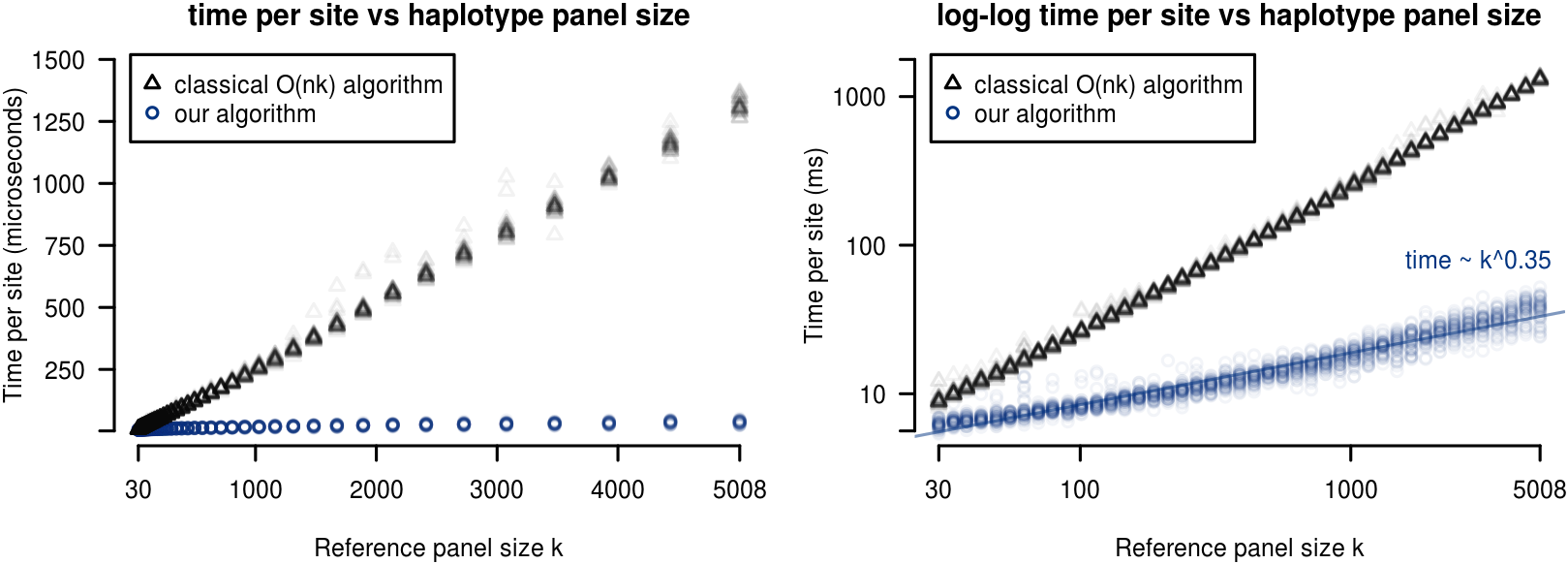
Runtime per site as a function of haplotype reference panel size k for our algorithm (blue) as compared to the classical linear time algorithm (black)

#### 5.3.1 Lazy evaluation of dynamic programming rows

In the average case, the time complexity of our lazy evaluation algorithm does not contribute to the overall time complexity of the algorithm. (Fig. 6, right) The lazy evaluation runtime also contributes minimally to the total runtime of our algorithm. (Fig. 6, left)

**Figure 6.**
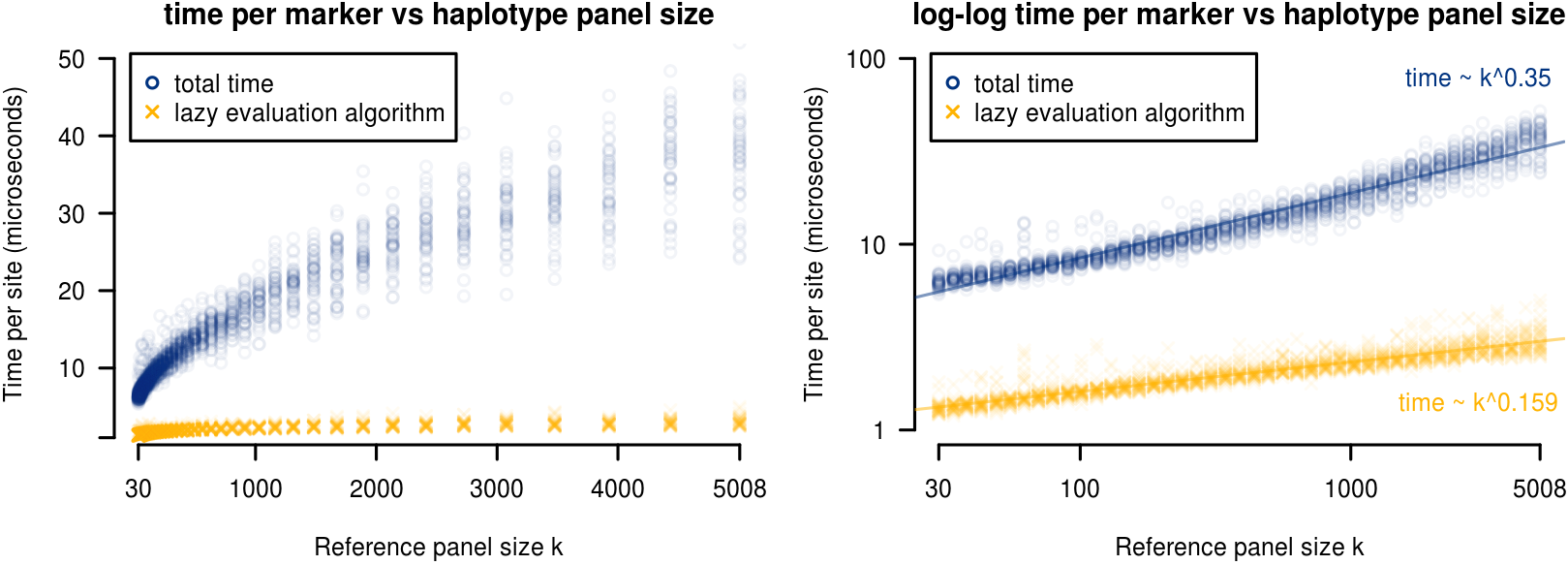
Time per site for the lazy evaluation subalgorithm (yellow) vs. the full algorithm (blue)

### 5.4 Sparse haplotype encoding

#### 5.4.1 Generating our sparse vectors

We generated the haplotype panel data structures from (§2) using the vcf-encoding tool vcf2slls which we provide. We built indices with multiallelic sites, which increases their time and memory profile relative to the results in (§5.2) but allows direct comparison to vcf records. Encoding of chromosome 22 was completed in 38 minutes on a single CPU core. Use of *M* CPU cores will reduce runtime proportional to *M*.

#### 5.4.2 Size of sparse haplotype index

In uncompressed form, our whole genome *.slls index for chromosome 22 of the 1000 genomes dataset was 285 MB in size versus 11 GB for the vcf record using `uint16_t`’s to encode haplotype ranks. When compressed with gzip, the same index was 67 MB in size versus 205 MB for the vcf record.

In the interest of speed (both for our algorithm and the 𝒪(*nk*) algorithm) our experiments loaded entire chromosome sparse matrices into memory and stored haplotype indices as `uint64_t`’s. This requires on the order of 1 GB memory for chromosome 22. For long chromosomes or larger reference panels on low memory machines, algorithm can operate on sequential chunks of the reference panel.

## 6 Discussion and significance

To the best of our knowledge, ours is the first forward algorithm for any haplotype model to attain sublinear time complexity with respect to reference panel size. Our algorithms could be incorporated into haplotype inference strategies by interfacing with our C++ library. This opens the potential for tools which are tractable on haplotype reference panels at the scale of current 100,000 to 1,000,000+ sample sequencing projects.

### 6.1 Applications which use individual forward probabilities

Our algorithm attains its runtime specifically for the problem of calculating the single overall probability *P*(*o*|*H*, *ρ*, *μ*) and does not compute all *nk* forward probabilities. We can prove that if *m* many specific forward probabilities are also required as output, and if the time complexity of our algorithm is 𝒪(Σ_*i*_│*ϕ*_*i*_│), then the time complexity of the algorithm which also returns the *m* forward probabilities is 𝒪(Σ_*i*_│*ϕ*_*i*_│+ *m*).

In general, haplotype phasing or genotype imputation tools use stochastic traceback or other similar sampling algorithms. The standard algorithm for stochastic traceback samples states from the full posterior distribution and therefore requires all forward probabilities. The algorithm output and lower bound of its speed is therefore 𝒪(*nk*). The same is true for many applications of the forward-backward algorithm.

There are two possible approaches which might allow runtime sublinear in *k* for these applications. Using stochastic traceback as an example, first is to devise an 𝒪(*f*(*m*)) sampling algorithm which uses *m* = *g*(*k*) forward probabilities such that 𝒪(*f* ○ *g*(*k*)) < 𝒪(*k*). The second is to succinctly represent forward probabilities such that nested sums of the *nk* forward probabilities can be queried from 𝒪(*ϕ*) < 𝒪(*nk*) data. This should be possible, perhaps using the positional Burrows-Wheeler transform [5] as in [11], since we have already devised a forward algorithm with this property for a different model in [13].

### 6.2 Generalizability of algorithm

The optimizations which we have made are not strictly specific to the monoploid Li and Stephens algorithm. Necessary conditions for our reduction in the time complexity of the recurrence relations are

▸ Condition 1. The number of distinct transition probabilities is bounded.
▸ Condition 2. The number of distinct emission probabilities is bounded. Favourable conditions for efficient time complexity of the lazy evaluation algorithm are
▸ Condition 1. The number of unique update maps added per step is bounded.
▸ Condition 2. The update map extension operation is composition of matrices of bounded size. This can be generalized to a broad algebraic class^5^ of update operations provided that they have bounded runtime.

The reduction in time complexity of the recurrence relations depends on the Markov property, however we hypothesize that the delayed evaluation needs only the semi-Markov property.

### 6.2.1 Other haplotype forward algorithms

Our optimizations are of immediate interest for other haplotype copying models. The following related algorithms have been explored without implementation.

▸ Example (Diploid Li and Stephens). We have yet to implement this model but expect average runtime at least subquadratic in reference panel size *k*. We build on the statement of the model and its optimizations in [9]. We have found the following recurrences which may be combined with a system of lazy evaluation algorithms:

#### ▸ Lemma 11

*The diploid Li and Stephens HMM may be expressed using the recurrences*

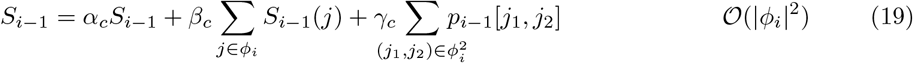

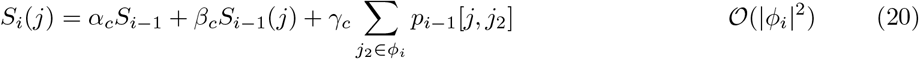

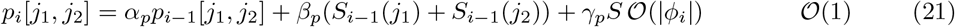

*where α*_(·)_,*α*_(·)_,*α*_(·)_ *depend only on the diploid genotype o*_*i*_.

▸ Example (Multipopulation Li and Stephens). [4] We maintain separate sparse haplotype panel representations 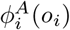 and 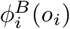 and separate lazy evaluation mechanisms for the two populations *A* and *B*. Expected runtime guarantees are similar.

This model, and versions for > 2 populations, will be important in large sequencing cohorts (such as NHLBI TOPMed) where assuming a single related population is unrealistic.

▸ Example (More detailed mutation model). It may also be desirable to model distinct mutation probabilities for different pairs of alleles at multiallelic sites. Runtime is worse than the biallelic model but remains average case sublinear.

▸ Example (Sequence graph Li and Stephens analogue). In [13] we described a hidden Markov model for a haplotype-copying with recombination but not mutation in the context of sequence graphs. Assuming we can decompose our graph into nested sites then we can achieve a fast forward algorithm with mutation.

▸ Example (Semi-Markovian recombination model). The lazy evaluation algorithm 3 may efficiently allow time-since-recombination dependent transition probabilities.

## Acknowledgements

We would like to thank Jordan Eizenga for his helpful discussions throughout the development of this work.

## Footnotes

1 Yohei Rosen was supported in part by a Howard Hughes Medical Institute Medical Research Fellowship.

2 We confirm these results in section 5.2.

3 The generalization is trivial

4 This approach is known as *lazy evaluation*.

5 Specifically, any collection of operations forming a *category* in the sense of category theory

